# Culturing primary neurons from rat hippocampus and cortex

**DOI:** 10.1101/491118

**Authors:** Madhusmita Priyadarshini Sahu, Outi Nikkilä, Seija Lågas, Sulo Kolehmainen, Eero Castrén

## Abstract

Primary neurons from rodent brain hippocampus and cortex has served as great tools in biomedical research over the years. However, protocols for the preparation of primary neurons vary, which often leads to conflicting results. This report provides a robust and reliable protocol for the production of primary neuronal cultures from the cortex and hippocampus with minimal contribution of non-neuronal cells. The neurons were grown in serum free media and maintained for several weeks without any additional feeder cells. The neuronal cultures maintained according to this protocol differentiate and by three weeks develop extensive axonal and dendritic branching. The cultures produced by this method show excellent reproducibility and can be used for histological, molecular and biochemical methods.

## Introduction

Primary culture of rodent hippocampal or cortical neurons remains one of the fundamental methods of modern neurobiology. Primary neurons can be easily collected and over a few days or weeks differentiate into a culture with clearly separable axons, dendrites, dendritic spines and synapses. By modifying culture medium and conditions, numerous factors directing different aspects of neuronal survival, differentiation and phenotype have been revealed.

Several more or less thoroughly described protocols for cultures of primary neurons have been published over years. Original methods used serum to support neuronal survival and differentiation (1) but more recently culture methods using defined media without serum have been introduced (2, 3, 4 2, 1490-8, 5). Glial cells provide critical support to cultured neurons (6 277, 1684-7, 7 468, 223-31, 8), and methods to keep excess glial proliferation in check or to prevent mixing between neurons and glia have been published (9 1, 2406-15).

One of the disadvantages of primary culture is that they do not divide in culture and need to be generated from embryonic or early postnatal brains every time. Moreover, successful dissection and preparation of cultures requires substantial skill and experience. Over several decades, cell lines have been discovered and created that mimic many or most of the features of primary neurons (10 5, 327-30, 11), and more recently, differentiated stem cells from rodents (12) or humans (13) have been introduced as alternatives for primary cultures, but none of these have replaced embryonic primary neurons from their position as a gold standard for neuronal cultures.

Neuronal cultures, however, vary vastly depending on source, age of derivation and culture conditions. Results obtained by a culture protocol used in one lab may not be reproducible in another lab, which adds to the ongoing discussion about reproducibility crisis. We have over more than a decade developed and refined a culture protocol of primary neurons derived from E17 rat embryos, which has been successfully used in several publications (14, 15). Here, we describe this protocol, necessary materials and methods as well as the characteristics of neurons derived through it in detail and share the protocol with other groups with the aim to promote reproducibility and rigor.

## Materials and Methods

### Animals

Pregnant female rats were obtained from Envigo (Harlan Labs,UK). The plug date of the female rats was marked E0. All embryos staged at E17-18 from the female rats were used in the experiments. Animals were kept in standard conditions (temperature 22°C, 12-hour light/dark cycle). Food and water were available ad libitum. All the experiments were performed according to institutional guidelines (University of Helsinki internal license number: KEK17-016)

### Composition of different solutions used

1. PBS buffer with pH= 7.4 consist of 80g NaCl, 0,2g KCl, 14g Na_2_HPO_4_ × 2H_2_O, 2g KH_2_PO_4_ in 1 liter Milli Q water. The final solution was autoclaved.
2. Preparation medium contains HBBS, 1mM sodium puruvate, 10 mM HEPES and the final ph 7, 2.
3. DMEM++ contained DMEM, 10% Fetal Bovine Serum, 1% L-glutamine, 1% penicillin-sreptomycin
4. Papain solution included 0.5 mg papain, 10 μg DNAse I in 5ml Papain Buffer. The stock Papain buffer comprised of 1mg DL-Cystein HCl, 1mg BSA, 25mg Glucose in 5ml PBS
5. Trituration medium consists of 10 μg Dnase I in 10ml Preparation medium as above.
6. Growing medium consists of Neurobasal medium, 2% B27 supplement, 1% L-glutamine, 1% penicillin-streptomycin
7. Poly-L-lysine working solution was made at 1:10 dilution Poly-L-lysine in MilliQ water
8. 4% PFA consists of 40 g paraformaldehyde in 1 litre PBS (1). The final solution was filtered.
9. PBST consists of 0.3% Triton X-100 in PBS
10. Blocking buffer consists of 1% BSA, 4% normal goat serum, 0.3% Triton-X100 in PBS
11. Primary antibodies used were GFAP, NeuN and MAP2 at 1:1000 dilution in the blocking buffer.
12. Secondary antibodies were Goat Anti Rabbit 647, Goat Anti Mouse 568, and Goat Anti Chicken 488 diluted 1:1000 in blocking buffer.

### Extraction procedures

Timed pregnant female rats were terminally anesthetized with carbon dioxide. Abdominal skin was washed with 70% ethanol followed by opening of the peritoneal cavity (Video file-1). Amniotic sacs were exposed with fine scissors and embryos were taken out from the uterus. These embryos were transferred to 50 ml tube with PBS on ice. The heads were immediately moved to ice-cold PBS. Brain dissection was performed in the preparation medium (2) on 60 mm petridish on ice. For precise identification of the brain structures, the cortex and hippocampus dissection were performed under the light microscope at room temperature. Step by step dissection has been compiled in a series of video files supplied as supplementary video files (Video files 2-5). The cortex tissue was let to recover in 10 ml DMEM (3) on a 90 mm petridish. The hippocampus was added into the preparation medium (2) on 35 mm petridish on ice. Cells were handled separately according to the following instructions. From here onwards, all the experiments were carried out under sterile conditions.

#### Cell preparation for hippocampus

1. Pre-coat multi-well plates (for biochemical assays) or round coverslips within a 4-well plate (for staining) overnight with Poly-L-lysine (7), 10 μg/ml at 37°C. Next day, before plating, wash the plates twice with PBS.
2. Transfer the dissected hippocampus tissue into pre-warmed papain solution (4) at +37°C.
3. Incubate the tissue for 5-10 min at +37°C. After the tissue sinks to the bottom, excess of papain solution is discarded.
4. Cells were triturated in about 3 ml trituration medium (5) with 20G needle with 10 ml syringe 10 times. Let the undissociated tissue sink to the bottom for about 30-60 sec. Transfer the supernatant to a fresh 15 ml tube.
5. Repeat trituration 2-3 times.
6. Centrifuge for 5 min at 900 rpm/154 g in Centrifuge 5810 (Eppendorf).
7. Remove the supernatant and resuspend the cell pellet in fresh growing medium (6). The hippocampal cells were diluted to 1:10 (5 μl +45 μl growing media). 10 μl of the diluted cells was plated into the disposable hemocytometer. Cell counting was done under a light microscope and using a manual cell counter.
8. For immunostaining, we typically use 50 000 cells per 1.9 cm^2^ coverslip within a 4-well plate. The cells were let to grow in +37°C (5% CO_2_). Dilute and plate at a density of 3 000-500 000 cells in different well format of 4, 12, 24 or 96 well plate, depending on the duration and purpose of the experiment.

#### Cell preparation for cortex

1. Pre-coat multi-well plates (for biochemical assays) or coverslips within a 4-well plate (for staining) overnight with Poly-L-lysine (7), 10 μg/ml at 37°C. Next day, before plating, wash the plates twice with PBS.
2. Triturate cortex in DMEM (3) with 20G needle with 10 ml syringe 12-15 times (Video file-6).
3. Centrifuge 5 min at 900 rpm/154g.
4. Re-suspend the cell pellet in 2ml/brain DMEM (3), let it sit down for 2-3 minutes to allow unwanted residue to sediment.
5. Pre-plate the supernatant on 90mm Petri dish. This is done to get clear of the glial cells by allowing them to attach to the bottom of the plate. Incubate the cells at +37°CO_2_, 5% COL for 30 min.
6. Carefully remove the supernatant, avoid taking the attached cells from the bottom of the plate.
7. Centrifuge 5 min at 900 rpm. Resuspend the pellet in growing medium (5). Let it sit down 2-3 min to sediment the debris.
8. Remove the cell suspension into a new tube.
9. Count cells using the 0.4% of Trypan Blue dye. The cortical cells were used at 1:20 dilution (5μl +75 μl growing medium +20μl trypan blue) for cell counting. The cell counting was done similarly as the hippocampal cells.
10. For immunostaining, we typically use 50 000 cells per 1.9 cm^2^ coverslip within a 4-well plate. The cells were let to grow in +37°C (5% CO_2_). Dilute and plate at a density of 3 000-500 000 cells in different well format of 4, 12, 24 or 96 well plate, depending on the duration and purpose of the experiment.

Both the cell types were grown for 7, 14 and 21 days at +37°C with 5% CO_2_ incubator. Cells were fixed at respective time points with 4% PFA (8) for 15 minutes followed by quick wash with PBS.

#### Immunohistochemistry

1. The 4 well plates were washed 3×5 min with PBST (9).
2. The fixed cells were blocked with blocking buffer (10) for 1 hour at room temperature.
3. Blocking buffer was washed away and primary antibody (11) prepared in the blocking buffer (9) was added. This was left on shaker at +4°C overnight.
4. The antibody was removed and washed 3×10 minutes with PBST.
5. After washing the secondary antibody prepared in the same blocking buffer was added to the wells at a dilution of 1:1000. The cells were left o/n on shaker at +4°C. The plates were covered with foil to protect from light.
6. The secondary antibody was washed with 1×10 minutes PBST followed by 2×10 minutes with PBS.
7. The coverslips were cleaned in MilliQ and mounted using mounting media on to the microscope slides.
8. The slides are stored in dark protected from light at +4°C until further use.

#### Imaging

The cells attached onto the coverslips were mounted on Superfrost slides (Thermo Scientific). The whole slide imaging was performed using Histoscanner (3D HISTEC Ltd., Hungary) at the genome biology unit, Biomedicum Helsinki. The images were analyzed using panoramic viewer software (3D HISTEC Ltd., Hungary). Four coverslips for each stage was scanned. Cell counting was performed using ImageJ software (https://imagej.nih.gov). Cells from a single focal plane was analyzed for an area of 4 mm^2^ per image per coverslip. Five images from each coverslip was counted and averaged.

The high magnification images were acquired using a Zeiss LSM 880 confocal microscope, at the Biomedicum Imaging unit, Biomedicum Helsinki. The lasers used were Alexa 488, Alexa 565, Alexa 647 and Alexa 405. To minimize crosstalk between different channels, the images were acquired by sequential scanning. The most commonly used algorithm for image acquisition was the maximum projection. The maximum intensity method is useful in extracting and detecting finer structures in a three-dimensional mode.

#### Statistical analysis

All the data was analyzed using the GraphPad Prism 6 software (La Jolla, CA, USA). The groups were compared using One-way ANOVA. All the data are represented as means ± SEM.

### Cell maintenance

Cells were grown at +37°C in incubator (Heraeus, model Heracell) with 5% CO_2_ gas. Half of the growing medium was changed once a week. The cell density was 50 000 cells in 1.9 cm^2^ area. The cells have been fixed at different time points at 7, 14 or 21 DIV.

## Results

The cell culture method described here has been developed and used in our lab for over a decade. The quality of neurons has been high and reproducible over the period of time. The overall procedure is outlined in the Figure 1. The extraction of the pups was performed under semi-sterile conditions within the animal facility (Fig 1A). The brain extraction and dissection were performed in a laminar hood under sterile conditions (Fig 1 B). The trituration was performed separately for hippocampal neurons and cortical neurons (Fig 1C). The quality of the cell dissociation was assessed by diluting and counting the cells using the hemocytometer (Fig 1D). The cortical cells were dyed with trypan blue to detect dead cells. The cortical cells along with the dye was used for cell counting (Fig 1Db).

**Figure 1:**
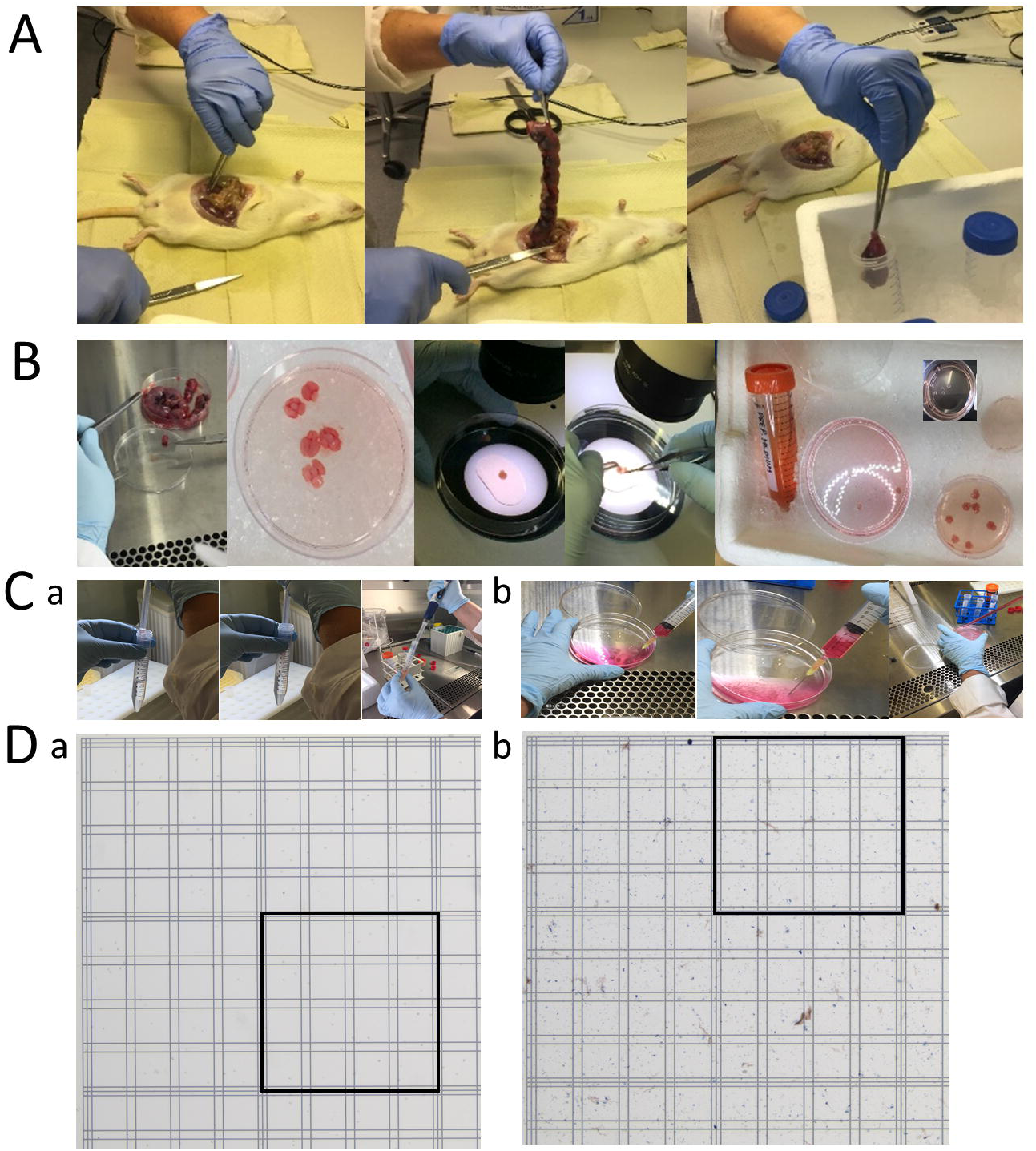
The procedure for extracting neuronal cells from the intact animal tissue. A. This part of the procedure was performed in the animal facility a-opening the visceral cavity of the rat, b-extracting the pups, c-collecting the pups into sterile PBS. B. This procedure was performed in the sterile laminar hood. a-c Extraction of the brain from the pups, d-e dissecting the cortex and hippocampus from the brain. C. Trituration of the tissue to produce homogenous cells, a-c hippocampal neurons and d-f cortical neurons. D. Cell counting using the Bürker slide a-hippocampal neurons, b-cortical neurons with trypan blue.

**Figure 2:**
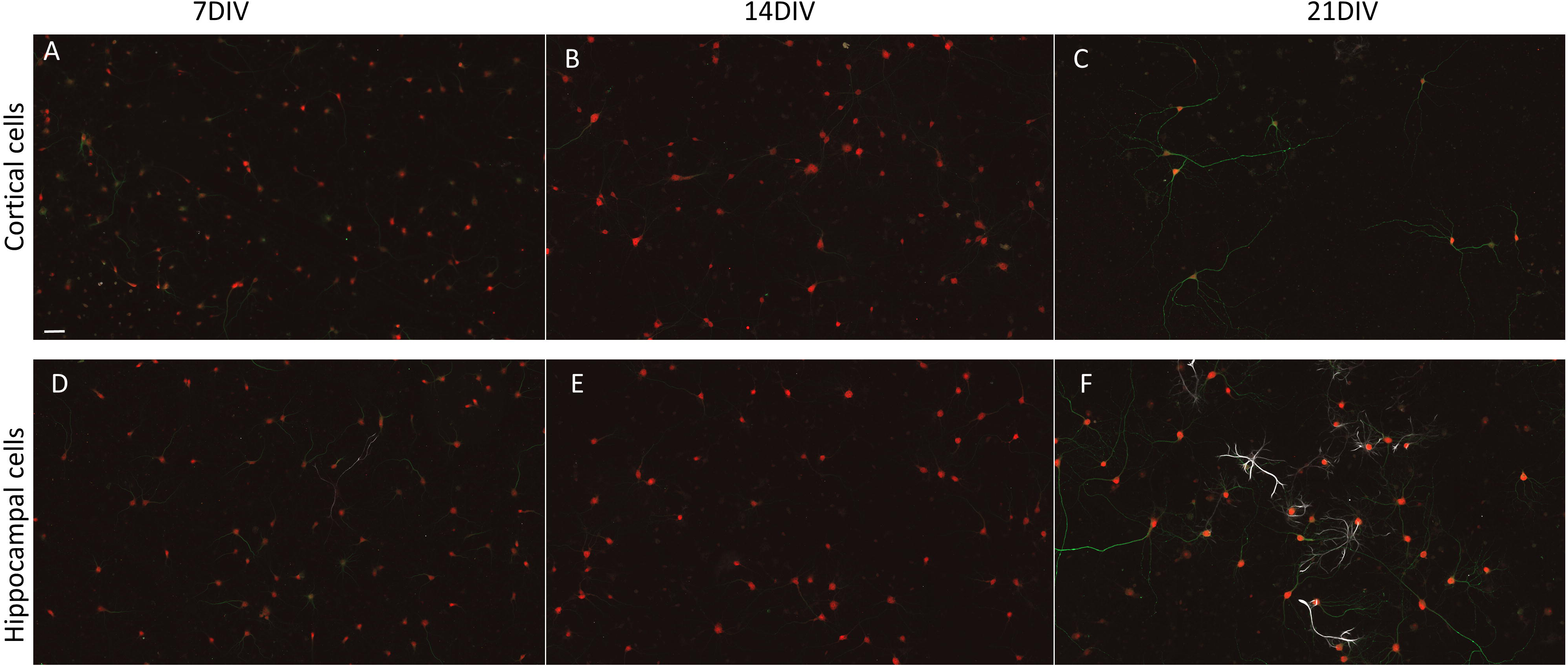
Culturing cortical and hippocampal neurons at different time points. A-C Cortical neurons at 7DIV (A), 14DIV (B) and 21DIV (C). D-F-Hippocampal neurons at 7DIV (D), 14DIV (E) and 21DIV (F). The neurons are stained with NeuN (red), Map2 (green) and GFAP (grey).

The markers for cell staining was selected to identify neurons from glial cells if any were present. The level of maturation of these cells was studied using the dendritic markers. The cells were stained for markers such as NeuN (stains neuronal nuclei), GFAP (stains glial cells) and Map2 (stains neuronal somato-dendritic compartment) at three different time points. The density of cells was about 50,000 cells for both hippocampal and cortical cells. In the cortical cells, no GFAP positive cells were detected at 7DIV and 14DIV. At 21DIV the GFAP cells formed 4% of the total cell population in the primary cortical cells (Fig-3a). The number of cortical neurons sharply decline at 21DIV. For primary hippocampal neurons, the number of GFAP positive cells were 2% at 7DIV, 6% at 14DIV and 28.5% at 21DIV of the total cell population (Fig-3b). The dendritic branching in both the cells increase from 7DIV until 21DIV (Fig - 4).

**Figure 3:**
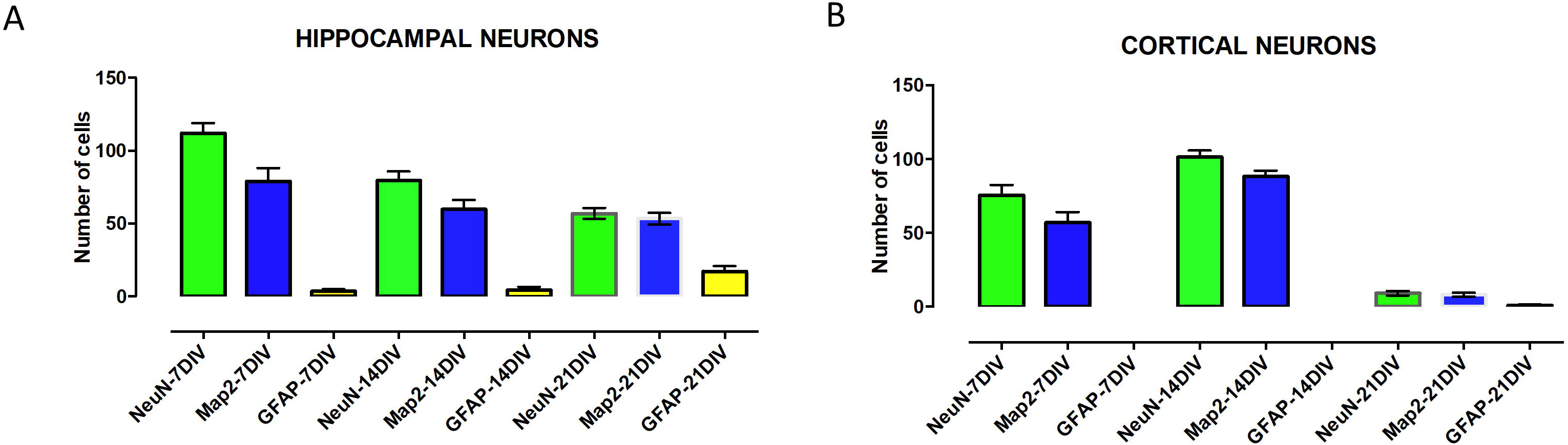
Cell counting in both hippocampal and cortical cells using markers for neurons (NeuN), astrocytes (GFAP) and a dendritic cell marker, Map2. The cell counting numbers of the representative markers in cortical neurons(A) and hippocampal neurons (B).

**Figure 4:**
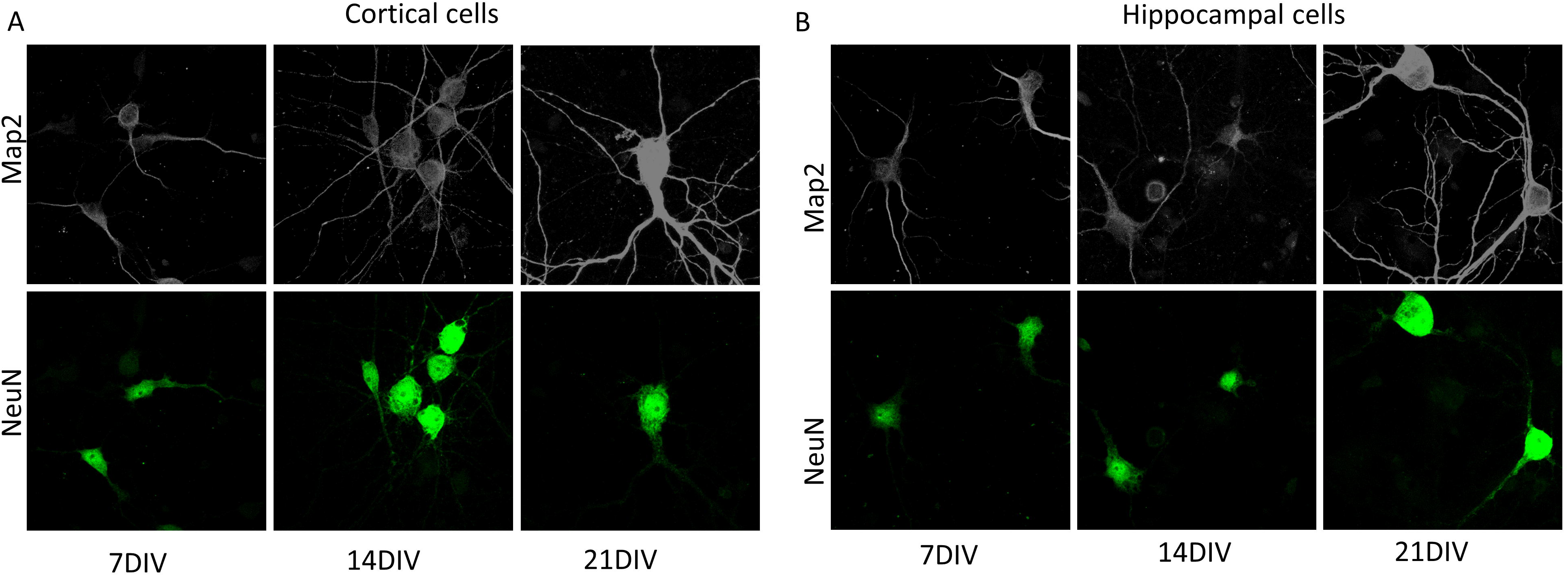
Higher magnification image of the cells stained for markers Map2 and NeuN. The Map2 branching increase at 14DIV and at 21DIV with more tertiary branches being observed both in the cortical cells (A) and hippocampal cells (B).

## Discussion

The primary neuronal cell culture is an ideal system for investigating cellular mechanisms at a higher resolution. The current protocol generates relatively pure neuronal cultures with maximum reproducibility and minimal contribution of glial cells. In this method we have cultured the neurons for 3 weeks without any additional feeder cells. For cell dissociation we have used papain only for hippocampal cells rather than trypsin, while cortical cells were dissociated by trituration without any prior enzymatic digestion. It has been observed that trypsin digestion of tissue leads to RNA degradation (16). The cortical cells recover from mechanical dissociation through a short incubation at 37 °C in DMEM with serum. The growth of non-desired cells, especially glial cells have been minimized in this study. This has been achieved by incubating at 37 °C and carefully removing the supernatant containing unadulterated cortical neurons. Low levels of glia are important especially if neurons will be used for biochemical analysis. However, it is well known that in the complete absence of glia, neurons fail to make efficient synapses (6), so a small percentage of glial cells is an advantage. These cells could be grown on coverslips for staining purposes. For biochemical methods, the wells were directly coated with poly-L-Lysine overnight. After washing away the poly-L-Lysine, cells were plated directly into it.

The cell lines have been the largest source for medical research in the past due to their immortal nature. These immortal cell lines have resulted with variable results arising after different passage times. This has been described as genetic drift as cells are passaged (17). The primary cells lack the immortality factor and therefore are the best *in vitro* models for biomedical research of the nervous system. These cells are genetically more stable than neuronal cell lines. The primary cells in culture maintain many crucial markers and functions as seen *in vivo*. Thus, they complement the *in vivo* experiments allowing for more controlled manipulation of cellular functions and processes. Once neurons are cultured, advanced molecular and biochemical study is easy to perform. For example, successful CRISPR-cas9 gene editing has been achieved using primary neuronal cultures (18). Furthermore, the cellular dynamics can be easily monitored through live imaging and electrophysiology. These features have established primary neurons as an essential tool for drug testing, with an additional advantage of reduced animal usage. However, variability in preparation methods reduces reproducibility of data, therefore, we hope that the method published here could be adopted by several research group for reliable and reproducible culturing of primary neurons.

## Abbreviations

E17: -Embryonic day 17
DIV: -Days *in vitro*
NeuN: -neuronal nuclei
GFAP: -Glial fibrillary acidic protein
Map2: -Microtubule Associated Protein 2

## Author contribution list

MS and EC designed the experiments and wrote the manuscript. MS, ON and SL performed the experiments. SK edited the video files.

## Acknowledgements

The authors do not declare any conflict of interest. This work was financially supported by ERC grant #322742 – iPLASTICITY, the Sigrid Juselius Foundation, and Academy of Finland grants #294710 and #307416 to EC.

## Figure legends

**Table 1:** Detailed protocol about materials

| ***Reagents*** | ***Equipments and Surgical instruments*** | ***Other materials*** |
| --- | --- | --- |
| NaCl (31434, Riedel) | Dissecting/stereo microscope (EZ4 HD, Leica) | Petridish 90mm (101RT/C, Thermo Scientific) |
| KCl (31248, Riedel) | CO<sub>2</sub> Incubator (Heracell, Heraeus) | Petridish 60mm (628102, Greiner) |
| Na<sub>2</sub>HPO<sub>4</sub> x 2H<sub>2</sub>O (0326, J.T. BAKER) | Centrifuge 5810 (Eppendorf) | Petridish 35mm (627102, Greiner) |
| KH<sub>2</sub>PO<sub>4</sub>(4871, Merck) | Scissors (RU 1003-14 Rudolf) | Multidish 4 well plate (176740, Thermo Scientific) |
| HBBS (14170088, Gibco) | Spring scissors (15004-08 Fine Science Tools) | Superfrost slides (J1800AMNT, Thermo Scientific) |
| 100mM sodium puruvate (11360039, Gibco) | Forceps (11000-13 Fine Science Tools) | Coverslips (Round 13mm, Thermo scientific) |
| 1M HEPES ph 7,2 (101926, ICN) | Curved forceps (11271-30/Dumont#7 Fine Science Tools) | C-Chip Disposable Hemocytometer (Bürker, LabTech) |
| DMEM (BE12-614F, BioWhittaker Lonza) | Straight forceps (11295-10/Dumont#5 Fine Science Tools) | Microscope slides ( ECN 631-1551, VWR) |
| Papain (P-4762, Sigma) |  |  |
| Dnase I (D4527, Sigma) |  |  |
| DL-Cystein HCl (C-9768, Sigma) |  |  |
| BSA (A7638, Sigma) |  |  |
| D-(+)-Glucose anhydrous (49139, Fluka) |  |  |
| PFA, Paraformaldehyde (1157, J.T. BAKER) |  |  |
| Poly-L-lysine (P4707, Sigma) |  |  |
| Normal goat serum (16210-064, Lifetechnologies, Gibco) |  |  |
| Triton-X100((93426, Fluka) |  |  |
| GFAP (12389, Cell Signaling Technology) |  |  |
| NeuN (MAB377X, EMD Millipore) |  |  |
| MAP2 (ab5392, Abcam) |  |  |
| Goat Anti Rabbit 647 (A21245, Life Technologies) |  |  |
| Goat Anti Mouse 568 (A11004, Life Technologies) |  |  |
| Goat Anti Chicken 488 (A11039, LifeTecnologies) |  |  |
| Mounting media with DAPI (ab104139, Abcam) |  |  |
| Trypan blue (T8154, Sigma) |  |  |

